# Phonological and temporal regularities lead to differential ERP effects in self- and externally generated speech

**DOI:** 10.1101/2021.05.04.442414

**Authors:** Alexandra K. Emmendorfer, Milene Bonte, Bernadette M. Jansma, Sonja A. Kotz

## Abstract

Some theories of predictive processing propose reduced sensory and neural responses to anticipated events. Support comes from M/EEG studies, showing reduced auditory N1 and P2 responses to self-compared to externally generated events, or when stimulus properties are more predictable (e.g. prototypical). The current study examined the sensitivity of N1 and P2 responses to statistical regularities of speech. We employed a motor-to-auditory paradigm comparing ERP responses to externally and self-generated pseudowords, varying in phonotactic probability and syllable stress. We expected to see N1 and P2 suppression for self-generated stimuli, with greater suppression effect for more predictable features such as high phonotactic probability and first syllable stress in pseudowords. We observe an interaction between phonotactic probability and condition on the N1 amplitude, with an enhanced effect of phonotactic probability in processing self-generated stimuli. However, the directionality of this effect was reversed compared to what was expected, namely a larger N1 amplitude for high probability items, possibly indicating a perceptual bias toward the more predictable item. We further observed an effect of syllable stress on the P2 amplitude, with greater amplitudes in response to first syllable stress items. The current results suggest that phonotactic probability plays an important role in processing self-generated speech, supporting feedforward models involved in speech production.

## 1 Introduction

The brain’s capacity to formulate predictions of upcoming events in our environment is one of the most studied phenomena across sensory modalities (e.g., Baldeweg, 2006; Blakemore et al., 2000; Rao & Ballard, 1999). These predictions may relate to the timing (‘when’, temporal prediction) and content/quality (‘what’, formal prediction) of upcoming sensory events (Arnal & Giraud, 2012; Kotz & Schwartze, 2010), and are based on our acquired knowledge and experience of the world. A special form of prediction generated by the brain is related to the sensory consequences of our own actions. The underlying mechanism is described by the internal forward model of motor control (e.g., Wolpert & Miall, 1996). According to this model, when a motor plan is formulated, an internal copy of the command, termed “efference copy” is used to generate a prediction of the anticipated sensory feedback. This prediction, or “corollary discharge”, is then compared to the actual sensory feedback (reafference signal), allowing the system to distinguish between self- and externally generated sensations, and to monitor and adapt our own motor output more readily. This model has also been applied to speech production, linking psycholinguistic models of feedback monitoring at the phoneme and syllable level, to general motor control mechanisms (e.g., Hickok, 2012; Kotz & Schwartze, 2016)

As a consequence of this mechanism, the sensory response to internally generated stimulation is suppressed, leading to well-known phenomena such as the inability to tickle oneself (Blakemore et al., 2000). This perceived sensory suppression, termed motor-induced suppression (MIS), goes hand in hand with the suppression of sensory-related neural activity, shown across multiple sensory domains, including somatosensory (Blakemore et al., 2000) and auditory (e.g., Christoffels et al., 2011; Knolle et al., 2012; Niziolek et al., 2013). The degree of MIS reflects the accuracy of the generated prediction: the better the match between predicted feedback and actual sensory feedback, the greater the suppression. This pattern is supported by observations of an inverse relationship between noise level and suppression in fMRI: with higher noise levels, less suppression is observed (Christoffels et al., 2011). The magnitude of the neural activity in response to self-generated sensations is therefore thought to reflect the prediction error, or the mismatch between predicted and actual feedback: Noisy situations result in less clear and less predictable feedback, leading to less suppression (i.e., more neural activity).

MIS is modulated by stimulus properties, including the predictability of the frequency and timing of tones (Bäss et al., 2008; Knolle et al., 2013a) or manipulations of voice identity (Johnson et al., 2021), voice quality and timing in speech (Aliu et al., 2009; Behroozmand & Larson, 2011). In a study comparing different utterances of the same vowel, self- or externally generated, Niziolek and colleagues (2013) observed greater suppression when the utterance showed formant ratios more prototypical for the individual speaker. Crucially, when the utterance deviated from the speaker’s prototype, the auditory cortical response predicted the correction of the articulation. This observation confirms that this mechanism may be involved in monitoring and correcting behavior. MIS is further modulated by experience, with musicians showing different suppression patterns than non-musicians (Ott & Jäncke, 2013). In summary, these findings suggest that greater suppression is indicative of more predictable sensory events, and that this suppression may be modulated by experience.

These observations suggest that MIS may be a suitable measure to investigate the brain’s sensitivity to regularities in the formal and temporal structure of speech during production. Within speech and language, regularities exist at multiple timescales, allowing the formulation of formal (e.g. phonotactic probability) and temporal (e.g. syllable stress) predictions across different processing levels. These predictions are established through exposure to regularities in speech throughout development, and evidence of sensitivity to these regularities is found already in infancy (Nazzi et al., 1998; Saffran et al., 1996). This sensitivity may provide an important foundation in the early stages of language acquisition, by allowing infants to segment the continuous speech signal into words (Jusczyk et al., 1999; Mattys & Jusczyk, 2001; Thiessen & Saffran, 2003), and continues to facilitate speech processing throughout the lifespan, as indicated by both behavioral and neural evidence.

Phonotactic probability modulates primarily sublexical language processes (i.e. independent of lexical/conceptual processing) in speech perception, such as nonword recognition (Luce & Large, 2001; Vitevitch & Luce, 1999) and recall (Thorn & Frankish, 2005). In contrast, lexical stress can guide the resolution of lexical conflict in spoken word recognition (e.g. <u>pre</u>sent = gift, pre<u>sent</u> = to give a presentation; Cutler, 2005). In production tasks, such as nonword repetition, both phonotactic probability (Edwards et al., 2004; Munson, Edwards, et al., 2005; Munson, Kurtz, et al., 2005; Vitevitch & Luce, 1998, 2005) and syllable stress (Vitevitch et al., 1997) modulate performance, with high phonotactic probability items and more frequently occurring stress patterns (e.g. in Dutch, first syllable stress) being repeated more accurately. Furthermore, lexical stress can guide the learning of novel phonotactic constraints (Bian & Dell, 2020).

There is ample neural evidence supporting the aforementioned behavioral observations during speech perception, with variations in phonotactic probability and stress patterns modulations neural processing (Bonte et al., 2005; Di Liberto et al., 2019; Emmendorfer et al., 2020; Rothermich et al., 2012; Tremblay et al., 2016). However, data on the neural correlates of these features in speech production is sparse. fMRI investigations have shown sensitivity to distributional statistics such as phonotactic probability, syllable frequencies or mutual information in speech production tasks across the speech network, including auditory as well as motor regions, with reduced BOLD signal for items with higher frequency of occurrence within the language (Papoutsi et al., 2009; Tremblay et al., 2016). These findings are in line with psycholinguistic models proposing that motor plans of more frequently occurring structures are stored in a “mental syllabary”, while less frequent articulatory representations need to be compiled from smaller units on the spot (Levelt, 1999; Levelt & Wheeldon, 1994; Schiller et al., 1996). Electrophysiological data on these features in speech production tasks is sparse. In a go/no-go task, where “go” decision was based on lexical stress position, N200 latency was earlier for words with first syllable stress (Schiller, 2006). However, this was proposed to be related to the incremental encoding (i.e. from word onset to end) of the meter during speech production, rather than a function of typical/atypical stress patterns, which is further supported by behavioral findings in trisyllabic stimuli (Schiller et al., 2006). Currently, we do not know of any studies investigating the effect of variations in phonotactic probabilities during speech production with electrophysiological methods.

The current experiment aimed to investigate how predictability of phonotactic probability and syllable stress contribute to speech production, extending our knowledge from previous studies investigating speech perception (e.g., Bonte et al., 2005; Emmendorfer et al., 2020) and production (e.g., Schiller, 2006; Tremblay et al., 2016). To approach this question, we focused our attention on motor-induced suppression, as this allows investigating how such (ir)regularities modulate the accuracy of the prediction generated through the efference copy. While some studies have investigated this phenomenon in overt speech production (e.g., Aliu et al., 2009; Christoffels et al., 2011; Niziolek et al., 2013), this comes with challenges due to artifacts caused from engaging the facial muscles during articulation. Furthermore, overt production leads to variability in the pronunciation of the individual utterances, which can lead to changes in the degree of suppression (Niziolek et al., 2013). This is a particularly relevant constraint in the current design, as less familiar features may show more variability in articulation as well as more speech errors (Heisler & Goffman, 2016; Munson, 2001; Sasisekaran et al., 2010). To circumvent these challenges, we employed a button-press paradigm, or motor-to-auditory paradigm, where the participant elicits the presentation of speech stimuli via button-press (e.g. Knolle et al., 2013a, 2019; Ott & Jäncke, 2013; Pinheiro et al., 2018).

The classical design in these experiments employs three conditions: an auditory-only condition (AO), where participants are passively presented with auditory stimuli, a motor-auditory (MA) condition, where participants trigger the generation of self-produced pseudowords through a button-press, and finally a motor-only (MO) control condition used to correct for the motor component (MA – MO = MAC). This design has been applied to investigate MIS in response to a range of stimulus types, including tones (Knolle et al., 2013a), voices (Pinheiro et al., 2018), vowels (Knolle et al., 2019), and single syllables (Ott & Jäncke, 2013). These designs typically elicit modulations of the auditory N1 and P2 components. Observed reduction of N1 amplitude in response to self-generated stimuli is thought to reflect an unconscious, automatic prediction resulting from the efference copy/corollary discharge, while P2 suppression reflects a more conscious differentiation between self- and externally generated events (e.g. Knolle et al., 2013a, 2019; Pinheiro et al., 2018). Here we investigated the effect of phonotactic and syllable stress regularities on MIS of the N1 and P2 components, using prerecorded utterances of bisyllabic Dutch pseudowords from each participant. Specifically, we aimed at testing the following hypotheses: (1) N1 and P2 amplitudes are reduced for self-generated stimuli compared to externally generated stimuli (i.e. main effect of condition, MIS), (2) this reduction in amplitude is modulated by phonotactic probability and syllable stress (i.e. interactions between phonotactic probability and condition, and syllable stress and condition), with high phonotactic probability and first syllable stress items leading to greater amplitude reduction due to greater predictability, and (3) phonotactic probability and syllable stress may interactively modulate motor-induced suppression (i.e. three-way interaction between phonotactic probability, syllable stress and condition), where we do not have precise predictions about the nature of this interaction.

## 2 Methods

### 2.1 Participants

34 right-handed native Dutch speakers participated in the study after giving their informed consent. The study was approved by the Ethical Committee of the Faculty of Psychology and Neuroscience at Maastricht University (ERCPN-OZL 205_17_03_2019) performed in accordance with the approved guidelines and the Declaration of Helsinki. Participants were invited to complete two sessions: one for recording the stimulus materials, followed by the EEG session. 5 participants completed the stimulus recording but did not complete the EEG session due to the COVID-19 pandemic. One participant was excluded from the EEG session due to failure to accurately reproduce stimuli. One participant was excluded due to excessive noise in the EEG signal (< 100 trials remaining per stimulus per condition). This led to a final sample of 27 participants (9 male, mean age: 21.9; standard deviation +/- 3.8), who completed both sessions of the experiment. The stimulus recording procedures and variations of the EEG paradigm were piloted in an additional 9 participants. The stimuli generated from these pilot participants were used to determine the criteria for stimulus selection as described in the following section.

### 2.2 Stimulus generation

The stimuli for the EEG experiment were prepared on an individual basis. Participants were invited for an initial stimulus recording session scheduled several days prior to the EEG session. The stimuli consisted of four pseudowords (Table 1), which differed from each other in phonotactic probability (notsal vs. notfal) and syllable stress (first vs. second syllable; adapted from Bonte et al., 2005; Emmendorfer et al., 2020). During the EEG experiment, each participant was presented with stimuli in their own voice. As second syllable stress is rare in Dutch, “natural” pronunciation of bisyllabic pseudowords with this stress pattern is challenging. To circumvent this issue, participants were presented with the target words, which were generated using a splicing procedure. The target words were spoken by a female Dutch speaker, who produced the syllables of interest by replacing them individually with syllables in existing bisyllabic Dutch words containing the same (spoken) consonant cluster and stress pattern as the target pseudowords (e.g. /**bad**zout/ → /**<u>not</u>**zout/ and /**bad**<u>sal</u>/ → **not**sal; /ont**slag**/ → /<u>not</u>**slag**/, and /ont**<u>sal</u>**/ → /not**sal**/; for more details see Emmendorfer et al., 2020). These spliced target words were presented to the participants of the current experiment. After ensuring the participant could hear and reproduce the differences between the pseudowords, each target was presented 15 times in random order, and the participants were asked to repeat them as accurately as possible. Participants were not explicitly instructed to attend to the stress pattern as this could lead to exaggerated expression of syllable stress.

**Table 1.**
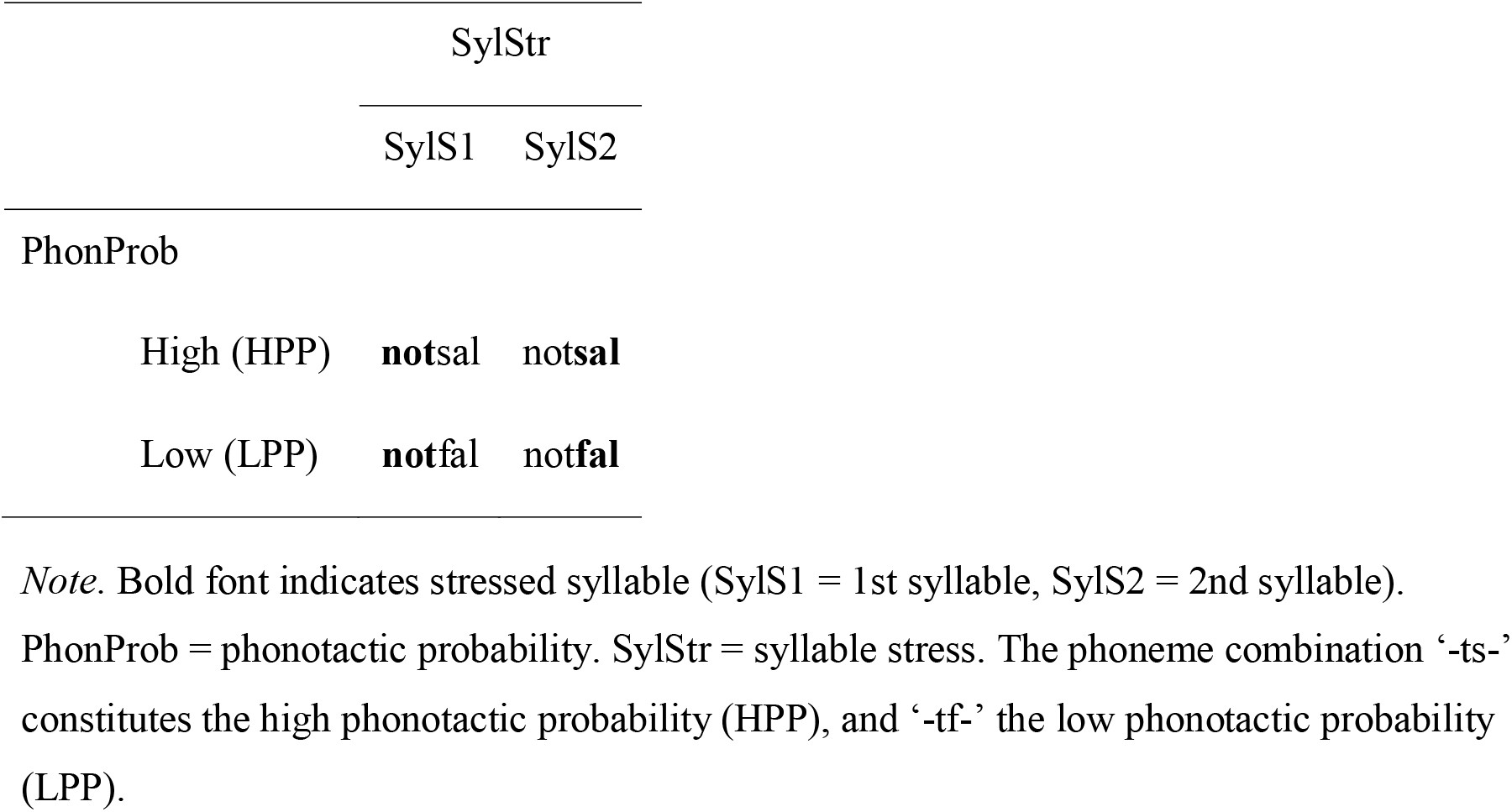
Stimuli

From the 15 repetitions of each pseudoword, one item was selected as the stimulus for the EEG experiment. To ensure comparability across participants, without having to manipulate the recording to deviate from the participants own naturally produced utterance, we selected items such that they were comparable in the timing of the perceptual centers (p-centers) of the syllables. P-centers are thought to represent the perceived “beat” of the speech stimulus. The timing of the p-centers was estimated with a beat detection algorithm (custom Matlab script adapted from Cummins & Port, 1998). Here, the beat, or p-center, is defined as the midpoint of each local rise in the amplitude envelope of the recorded signal, representing the vocalic nucleus of a syllable. The duration of the interval between the p-centers of each syllable in the bisyllabic pseudowords was calculated, and from 10 participants (9 pilot participants and 1 from the final sample), the average interval was calculated for each pseudoword. These values were used to select the best fitting stimulus for the participants who completed the subsequent EEG session. For each pseudoword, the item with the closest matching interval was selected. If this item contained acoustic artifacts or a mispronunciation, it was discarded, and the next best item was selected. This procedure allowed the selection of temporally comparable stimuli, while preserving each participant’s own pronunciation without editing or manipulating the timing. A representation of the stimuli included in the experiment can be found in Figure 1. Stimuli were filtered with a Hann bandpass filter (80 – 10500 Hz), and intensity scaled to 60 dB. Mean stimulus duration was 0.640 s (standard deviation: 0.056 s), and the mean interval between p-centers of the stimuli was 0.319 s (standard deviation: 0.042 s).

**Figure 1:**
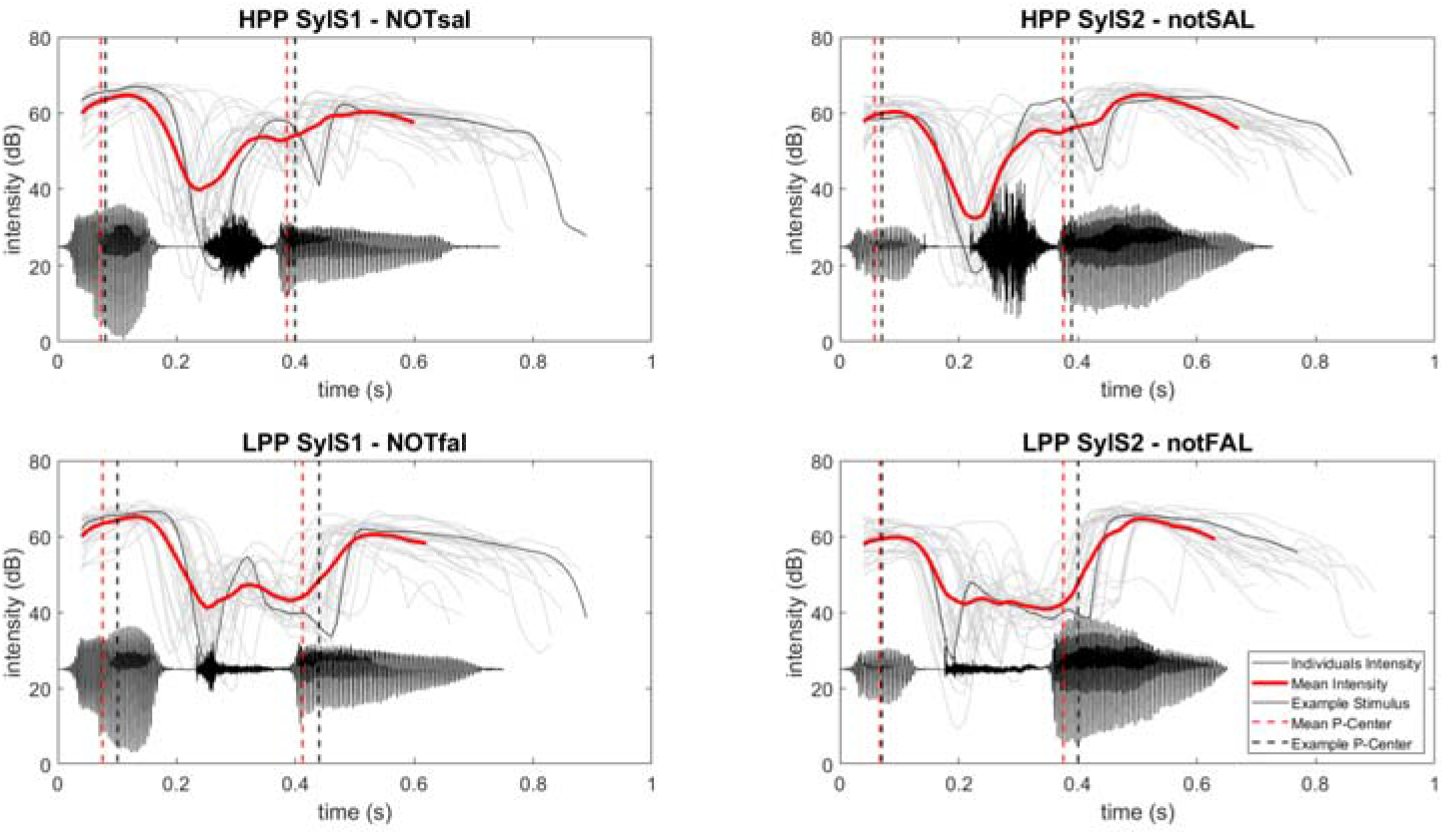
Stimuli. Stimuli selected for the EEG experiment. Individual intensity contours of the stimuli are represented in grey, mean intensity contours across participants in red. Stimuli from an exemplary participant are represented in black. Timing of the p-centers, representing the onset of the vocalic nucleus, are represented by dashed lines (red: averaged across participants, black: exemplary participant)

### 2.3 EEG paradigm

The paradigm (adapted from Johnson et al., 2021; Ott & Jäncke, 2013) consisted of three conditions (Figure 2A). In all three conditions, the trial began with the presentation of a fixation cross, followed by a cue (< left, > right) at 0.4 – 1.0 s after trial onset. In the motor-auditory condition (MA), participants pressed a button (left or right), which triggered the presentation of a stimulus. In the auditory-only condition (AO), participants were presented with the same cue, but the stimulus presentation occurred without button press, 0.5 s after cue onset. In the motor-only condition (MO), the participants pressed the cued button, but no stimulus was presented. This condition was included to correct for the motor component in the MA condition. This corrected motor-auditory condition (MAC) was calculated as MA – MO, thus allowing the comparison of neural activity in response to self-generated (MAC) and externally generated auditory stimuli (AO). A reduction in N1 and P2 amplitudes for MAC relative to AO is then interpreted as motor-induced suppression.

**Figure 2:**
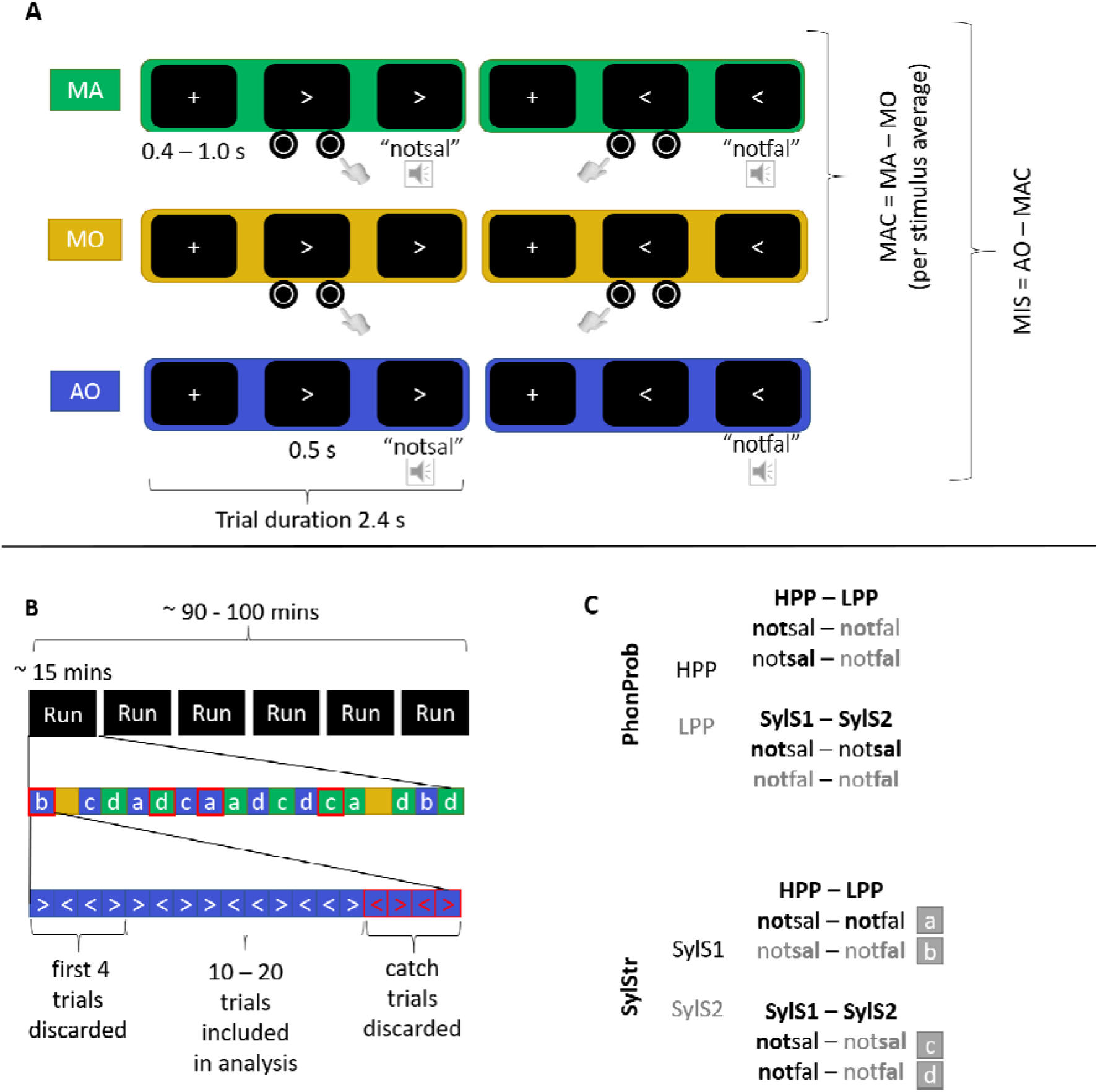
Experimental design. (A) Three experimental conditions: MA = motor-auditory, MO = motor only, AO = auditory only. (B) Overview of the EEG paradigm timeline. Letters a-d correspond to the stimulus pair presented as denoted in panel C. (C) Overview of stimuli and contrasted features: PhonProb = phonotactic probability, HPP = high phonotactic probability, LPP = low phonotactic probability, SylStr = syllable stress, SylS1 = first syllable stress, SylS2 = second syllable stress.

The EEG recording occurred over the course of 6 experimental runs, each consisting of 18 blocks (8 MA, 8 AO, 2 MO) (Figure 2B). In each MA and AO block, one stimulus pair was presented. The stimuli within the pair differed from each other in either phonotactic probability or syllable stress (Figure 2C), and each cue/button press corresponded to one stimulus. Each pair was presented twice per run and condition, with the cue/button assignment counterbalanced across blocks. Within each block, the first 4 trials (always including 2 left, 2 right) were excluded from analysis to allow the participant to form an association between cue and word. In four blocks per run (2 MA, 2 AO), four catch trials were included at the end of the block, where the cue-stimulus pairing was switched, i.e., the left cue was followed by the stimulus previously associated with the right cue. Participants were instructed to attend to the cue-stimulus pairing and were asked to report at the end of each block whether they noticed a switch. This task was included to ensure the participants were correctly associating the presented stimulus with the cue/button-press, and these trials were excluded from analysis. The total number of trials per block varied between 14 and 28 trials such that the participant could not anticipate when the catch trials would occur by counting. This resulted in 10 – 20 trials per block, and a total of 90 trials per condition/stimulus/cue assignment included in the analysis (Figure 2B).

### 2.4 EEG recording

EEG was recorded with BrainVision Recorder (Brain Products, Munich, Germany) using a 63-channel recording setup. Ag/AgCl sintered electrodes were mounted according to the 10% equidistant system, including 57 scalp electrodes, left and right mastoids for offline re-referencing, and four EOG electrodes to facilitate removal of artefacts caused by eye movements (2 placed on the outer canthi, 2 above and below the right eye). The scalp was cleaned at electrode sites and electrodes were filled with electrolyte gel to keep impedances below 10kΩ. Data was acquired with a sampling rate of 1000Hz, using Fpz as an online reference and AFz as ground. During recording, participants were seated on a comfortable chair in an acoustically and electrically shielded room.

### 2.5 EEG processing

EEG data was processed using the EEGLAB toolbox (Delorme & Makeig, 2004) and custom MATLAB scripts (MATLAB, 2018) The continuous EEG data were filtered using a bandpass filter of 1 – 30 Hz, and then downsampled to a sampling rate of 250 Hz. Noisy channels were identified, removed and interpolated using the EEGLAB plugin clean_rawdata, and the data were re-referenced to the average signal of the two mastoid electrodes. The data were then epoched 0 – 2.4 s relative to the onset of the trial to remove noisy break intervals, while still including the entire duration of the experimental blocks. The data were then decomposed using ICA. 2 – 4 independent components, reflecting blinks and horizontal eye-movements were removed for each participant. The reconstructed data were then baseline corrected to a window 0.2 s prior to the onset of the *cue*, and epoched -0.6 – 0.5 s relative the onset of the *stimulus* or *button-press* (this includes 0.1 s of the baseline window in the AO condition). This baseline window prior to the *cue* rather than the *stimulus* onset was selected due to an observed positivity in the AO response (Figure 3A), which was removed during the MA – MO subtraction. This deflection in the AO condition could not be removed from the data through highpass filtering or ICA, and likely reflects processes related to the visual cue (see Supplementary Figures S1, S2 and S3). This observation violates the assumption of baseline correction that there are no systematic differences across conditions in the selected window. Therefore, the pre-cue baseline window was deemed more appropriate. This notion is also supported by previous findings showing differences between self- and externally generated auditory stimuli already in the pre-stimulus window (Reznik et al., 2018). The pre-cue baseline correction is used throughout the analysis steps.

**Figure 3.**
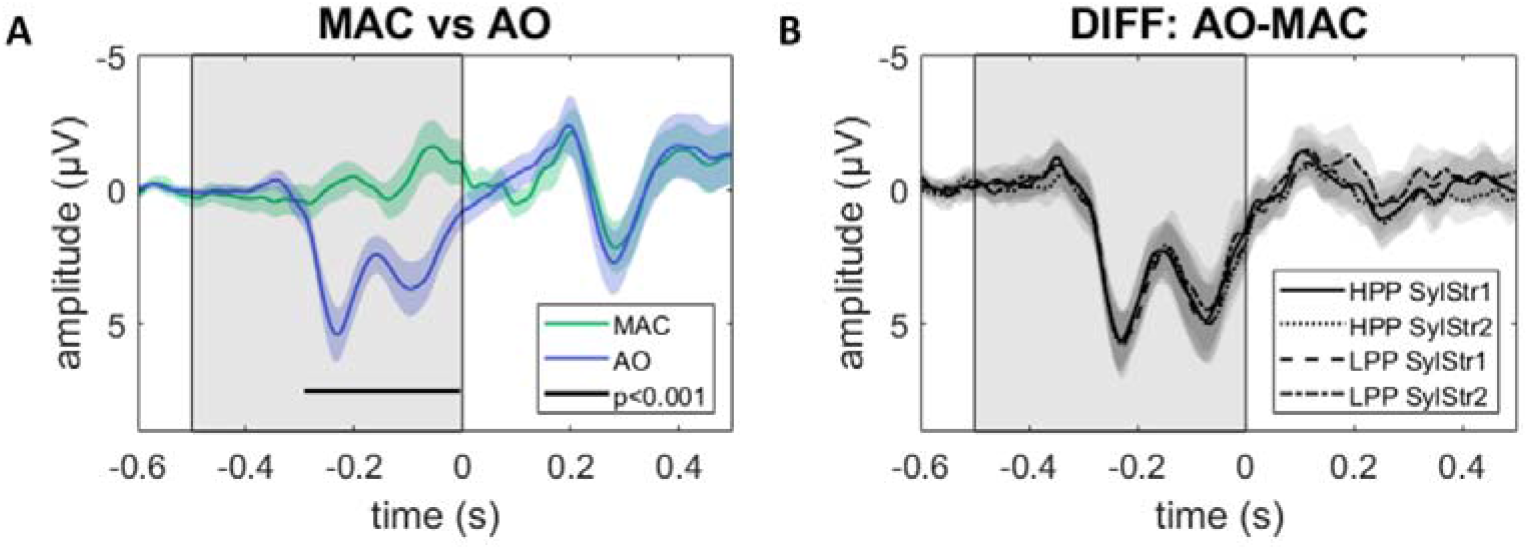
Overview of pre-stimulus deflection. (A) MAC (green) and AO (blue) conditions, time-locked to stimulus onset and averaged across stimuli (+/-95% CI of the mean) in a frontocentral ROI (FCz, FC1, FC2, FC3, FC4). The shaded window indicates the window where cluster-based permutation tests were performed. The black line indicates the timing of the observed cluster with a significant difference between MAC and AO. (B) AO – MAC difference wave for individual stimuli at the same frontocentral ROI. The shaded area indicates the window where cluster-based permutation tests were performed to test for systematic effects of PhonProb, SylStr or an interaction thereof. No significant differences were found.

Individual N1 and P2 peaks were manually determined from single subject average waveforms at electrode FCz for each stimulus and condition separately. While the auditory N1 and P2 are classically measured over the vertex electrode (Cz), we opted for a slightly more frontal site as visual inspection of the ERPs suggested the amplitudes to be less influenced by the pre-stimulus deflection at this channel (see Supplementary Figure S2). The N1 peak was determined as a negativity in the time window 100 – 300 ms following stimulus onset, P2 as a positivity following the N1 peak up until 400 ms. These time windows are later than the classically observed N1 and P2 windows, however, a relative delay is consistent with the nature of the stimuli due to their complexity (Conde et al., 2018) and slow onset rise time (Onishi & Davis, 1968). Furthermore, broad time windows were selected to determine the individual peak as we anticipated variability in their timing due to the variability of the individual stimuli (i.e., variations in rise time of first syllable between participants and between first and second syllable stress). If there was ambiguity in the selection of the peak within a waveform (e.g., 2 peaks within the given window), the peak with the more appropriate topography and timing relative to the participant’s average as well as the grand average across participants was selected, to ensure the inclusion of comparable neural events across conditions and participants. The amplitude in a window +/-24 ms surrounding this latency was extracted for all scalp electrodes.

### 2.6 Statistical analyses

Due to the pre-stimulus deflection observed in the AO condition (Figure 3A), we first investigated whether this indeed reflected a systematic difference between AO and MAC. Such a systematic difference between conditions would render a direct comparison of the N1 and P2 amplitudes of these two conditions invalid, as we cannot exclude that any observed modulations of these components might be driven by this deflection rather than true motor-induced suppression as hypothesized. We tested this via a cluster-based permutation analysis (Maris & Oostenveld, 2007). One-sided paired-samples t-tests between AO and MAC were performed at each time-point in the time-window -0.5 - 0 s relative to stimulus onset for 1000 random partitions using the ft_timelockstatistics function of the Fieldtrip toolbox (Oostenveld et al., 2011). This analysis revealed a significant difference between AO and MAC. The observed cluster started at approximately 0.3 s prior to stimulus onset, with a broad topographic distribution. Based on this unexpected observation, a direct comparison of AO vs MAC at N1 and P2 components could not be interpreted as motor-induced suppression. Due to this finding, we were unable to test our specific hypotheses regarding the modulation of motor-induced suppression by phonotactic probability and syllable stress. However, as the broader aim of this research was to investigate the role of these regularities in speech production, we pursue analyses to answer the question of how phonotactic probability and syllable stress modulate speech processing, and whether this differed across self- and externally generated speech.

Statistical analyses on N1 and P2 amplitudes were performed in R version 3.6.3 (R Core Team, 2013) using the rstatix package (Kassambara, 2019). Normal distribution of the N1 and P2 mean amplitude values was confirmed for all conditions via Shapiro-Wilk test (Supplementary Tables S1, S4), and outlier identification via boxplot methods did not reveal any extreme outliers (points beyond Q1 – 3*IQR, Q3 + 3*IQR). In a 2×2×2 (high vs. low PhonProb x first vs. second SylStr x AO vs. MAC Cond) repeated-measures ANOVA, we tested the following hypotheses for both N1 and P2 mean amplitudes averaged across electrodes in a frontocentral ROI (FCz, FC1, FC2, FC3, FC4): The N1/P2 amplitudes are modulated by the predictability of stimulus features (1) PhonProb and (2) SylStr, where more predictable utterances (i.e., high phonotactic probability and first syllable stress) lead to smaller amplitudes. Furthermore, (3) these features may interactively modulate N1/P2 amplitudes (PhonProb x SylStr interaction), and (4) may differ across conditions (PhonProb x Cond or SylStr x Cond interaction), where we would expect the MAC condition to show greater effects of these features due to error-monitoring. ANOVA results were corrected for multiple comparisons with Bonferroni-Holm correction using the adjust_pvalue() function (Cramer et al., 2016), and follow-up t-tests of simple effects were Bonferroni corrected.

## 3 Results

Visual inspection of the ERP grand averages (Figure 4A, 5A) reveals an N1/P2 morphology, with the N1 peaking around 200 ms and the P2 around 300 ms. When adjusted for the timing of the p-center of the first syllable of participants’ pseudoword pronunciations, the N1 and P2 latencies are shorter, at approximately 125 and 212 ms, respectively. For our analyses we kept the time-locking to stimulus onset as it resulted in delayed but better aligned N1 and P2 responses across participants. In the following sections, we present the results of the statistical analyses. Here, we report only significant or otherwise noteworthy main effects and interactions, as well as post-hoc simple effects. The full results of the statistical analyses can be found in the supplementary materials (Supplementary Tables S1, S2 and S3 for N1, Tables S4 and S5 for P2 results).

**Figure 4:**
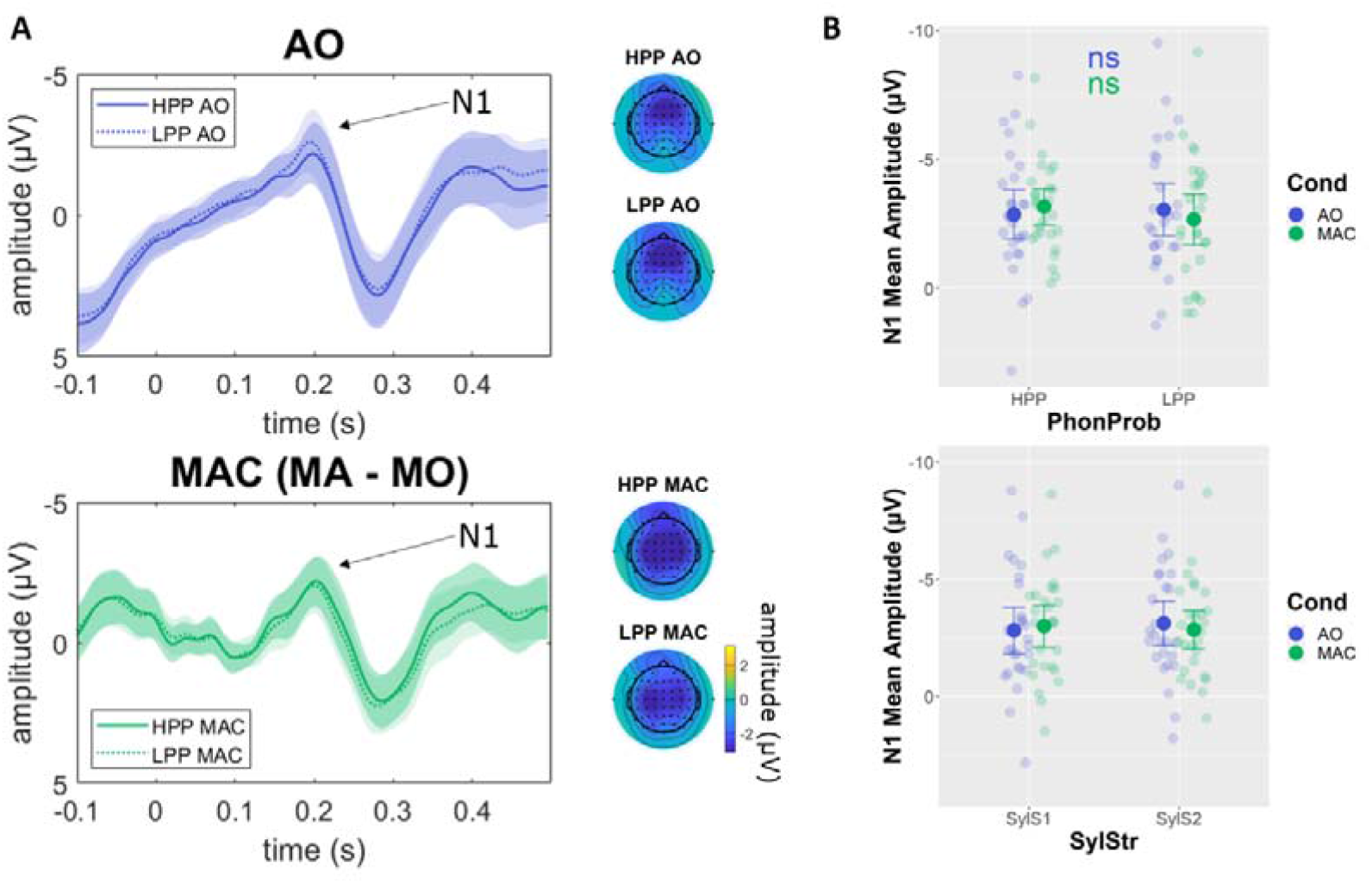
Interaction between phonotactic probability and condition on N1 amplitude. (A) ERP waveforms at frontocentral ROI (FCz, FC1, FC2, FC3, FC4) time-locked to stimulus onset, and corresponding topographies of N1 mean amplitudes at frontocentral ROI. (B) N1 mean amplitudes (+/- 24 ms surrounding individual peaks). HPP = high phonotactic probability, LPP = low phonotactic probability, AO = auditory only, MA = motor-auditory, MO = motor only, MAC = motor-auditory corrected, PhonProb = phonotactic probability, Cond = condition, ns = non-significant. Note: simple effects were not tested for SylStr x Cond as this interaction was not significant.

### 3.1 N1

A 2×2×2 repeated measures ANOVA (high vs. low PhonProb x first vs. second SylStr x AO vs. MAC Cond) on N1 mean amplitudes (+/- 24 ms surrounding peak) averaged within the frontocentral ROI revealed a significant interaction between phonotactic probability and condition (F(1,26) = 8.463, p.adj = 0.049, η^2^_P_ = 0.246). In the AO condition, LPP stimuli had a slightly larger N1 mean amplitude compared to HPP stimuli, while the reverse directionality was observed in the MAC condition (Figure 4A, B). This interaction was resolved by means of post-hoc paired samples t-tests testing the effect of phonotactic probability at each level of condition (Bonferroni corrected), which showed no significant effects for either AO (t(26) = 0.840, p.adj = 0.818, d = 0.162) or MAC (t(26) = -2.04, p.adj = 0.104, d = -0.392). Thus, the observed effect seems to reflect a crossover interaction, where the difference between HPP and LPP is different across conditions, but in neither AO or MAC do HPP and LPP differ from each other significantly. However, the effect size and mean amplitude difference is larger for MAC (d = -0.392, N1 mean amplitude HPP = -3.17 µV vs LPP = -2.67 µV) than AO (d = 0.162, N1 mean amplitude HPP = -2.87 µV vs LPP = -3.05 µV). No other main effects or interactions on N1 mean amplitude were significant (Figure 4B). We note that there was a small but significant difference in N1 latency across conditions (t(26) = -2.43, p = 0.022, d = -0.468), with the N1 peaking slightly earlier in the AO condition (mean latency = 190 ms) compared to MAC (mean latency = 197 ms).

### 3.2 P2

A 2×2×2 repeated measures ANOVA (high vs. low PhonProb x first vs. second SylStr x AO vs. MAC Cond) on P2 mean amplitude (+/-24 ms surrounding peak) averaged within the frontocentral ROI revealed a significant main effect of syllable stress (F(1,26) = 22.993, p.adj < 0.001, η^2^_P_ = 0.469). Stimuli with first syllable stress elicit a larger P2 (mean amplitude = 3.30 µV) compared to those with second syllable stress (mean amplitude = 2.39 µV; Figure 5). No other main effects or interactions were significant.

**Figure 5:**
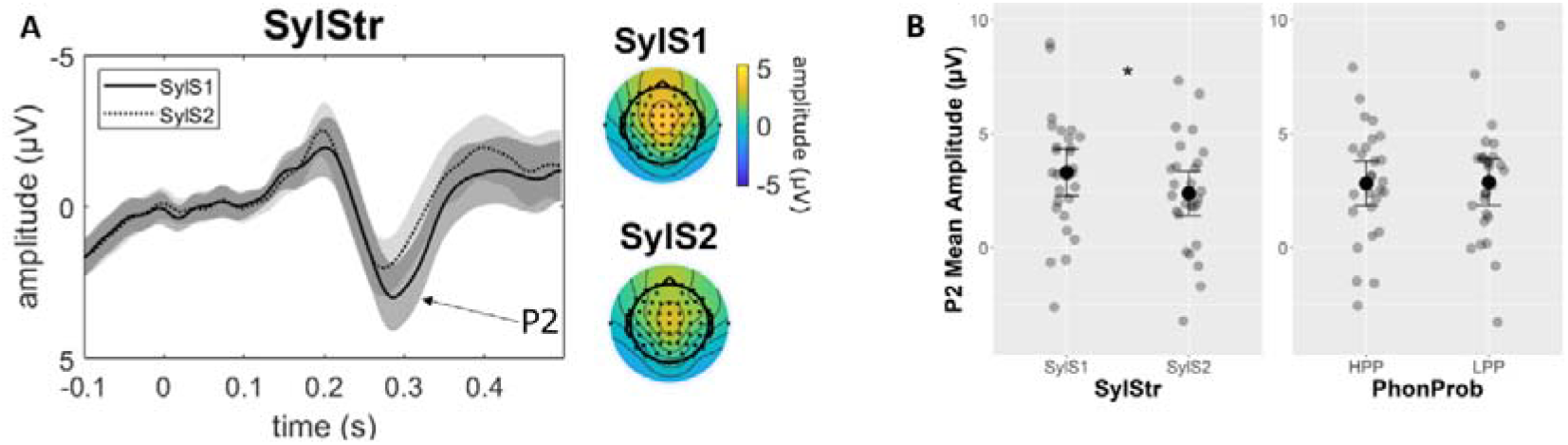
Main effect of syllable stress on P2 amplitude. (A) ERP waveforms at frontocentral ROI (FCz, FC1, FC2, FC3, FC4) time-locked to stimulus onset, averaged across phonotactic probability and condition, and corresponding topographies of individual P2 mean amplitudes (+/- 24 ms surrounding individual peaks). (B) P2 mean amplitudes (+/- 24 ms surrounding individual peaks). * p < 0.001. SylStr = syllable stress, SylS1 = first syllable stress, SylS2 = second syllable stress, PhonProb = phonotactic probability, HPP = high phonotactic probability, LPP = low phonotactic probability.

## 4 Discussion

The current study aimed to investigate whether motor-induced suppression of the N1 and P2 amplitudes is modulated by formal (phonotactic probability) and temporal (syllable stress) predictability in the speech signal. We used a motor-to-auditory paradigm, where participants triggered the generation of self-produced pseudowords through a button-press. This approach was intended as a step towards investigating speech production, while limiting the interference of motor artifacts and speech errors present during overt production of pseudowords. We expected to observe a motor-induced suppression effect, with larger N1 and P2 amplitudes in the auditory-only condition, compared to the motor-auditory condition. Furthermore, we expected this suppression effect to be modulated by phonotactic probability and/or syllable stress, where high probability items (high phonotactic probability and first syllable stress) would elicit greater suppression, as they might be more “prototypical” items in the language (Niziolek et al., 2013). Due to an observed pre-stimulus deflection in the auditory-only condition, not present in the motor-auditory condition after correcting for motor output (Figure 3), we were not able to test our specific a priori hypotheses regarding motor-induced suppression in the current data. However, the design still allowed investigating the broader question as to how phonotactic probability and syllable stress contribute to the processing of self- and externally generated speech. We observed a modulation of N1 amplitude by phonotactic probability, which was enhanced in response to self-generated stimuli, and a modulation of P2 amplitude by syllable stress, with first syllable stress eliciting a larger P2 compared to second syllable stress. While these analyses are post-hoc in nature, they provide insights into the differences in processing phonotactic and temporal regularities in self- and externally produced speech that can be followed up upon in future experimental designs.

### N1

We investigated the effect of variations of phonotactic probability and syllable stress across self- and externally generated conditions on N1 mean amplitude. Here, we observed a significant interaction between phonotactic probability and condition, but no significant main effects or simple effects. Inspecting the data revealed that this interaction was a crossover effect, indicating that the direction of the effect of phonotactic probability differs across conditions (i.e. HPP > LPP in MAC, LPP > HPP in AO). However, the simple effects did not reach significance in either the auditory-only or motor-auditory condition. It is noteworthy though that visual inspection of amplitudes as well as comparison of effect sizes revealed the difference between high and low phonotactic probability to be larger for self-generated words compared to externally generated ones. Thus, the pattern we observed in this interaction is in line with the notion that such regularities have greater weight in speech production due to feedforward processes. Variations in phonotactic probability of planned utterances may require different degrees of monitoring, as they may be more or less likely to result in mispronunciation. Indeed, it has been shown that phonotactic probability modulates accuracy and speed in speech production (Edwards et al., 2004; Munson, Edwards, et al., 2005; Munson, Kurtz, et al., 2005; Vitevitch & Luce, 2005). However, we hypothesized a larger amplitude for the less probable item as this would generate greater surprise, while our data suggest the opposite pattern. One could assume that these effects might be driven by differences in the motor-only (MO) condition, however we can exclude this possibility, as this condition does not include variations in stimulus type (i.e. same trials of MO are subtracted from all MA averages).

A closer look at theories of predictive processing may explain this discrepancy. While cancellation theories predict suppression of predicted sensory events to highlight novel or unexpected events (e.g., Blakemore et al., 2000), Bayesian theories suggest a perceptual bias or gain for predictable events (e.g., de Lange et al., 2018). Recent developments propose a two-process model to resolve the conflict between these contradictory theories (Press et al., 2020). Here it is proposed that the perceptual system is tuned toward expected events when there is a large overlap between prior and posterior probabilities (i.e. low surprise), resulting in perceptual gain for expected events, as suggested by Bayesian accounts (e.g., Thomas et al., 2020). When there is little overlap between these prior and posterior distributions (i.e. high surprise), this suggests that the model of the environment must be updated, resulting in higher activation to signal the unexpected event, in line with cancellation theories.

The pseudowords used in our design all consist of legal phonotactic structures and lexical stress patterns. Furthermore, the trials included in the analysis did not include any violations of predicted stimuli. Thus, the surprise generated by any given stimulus was low, and would not require the system to update their model of the world. Instead, given the acoustic similarity of high and low phonotactic stimuli, it is more likely that perception was biased toward the high probability item *notsal*. The trend toward a larger N1 amplitude for high probability items in the motor-auditory condition supports this notion. Interestingly, the timing of this modulation around 200 ms suggests that it may occur prior to, or concurrently with, the actual manipulation of phonotactic probability, which occurs at the syllable boundary (occurring around 200 – 250 ms, see Figure 1). This is in line with the opposing process theory proposed by Press and colleagues, as effects of perceptual sharpening are often observed prior to or within 50 ms of the expected stimulus, preceding cancellation effects (e.g., Press & Yon, 2019; Yon & Press, 2017). The perceptual sharpening may also render the system more sensitive to coarticulatory cues already present within the first syllable.

We did not observe a comparable modulation of the N1 by syllable stress, despite this feature also varying in probability in the Dutch language, with first syllable stress being the more common pattern in bisyllabic words. This observation is in line with a previous study in speech perception (Emmendorfer et al., 2020), where variations in syllable stress did not modulate MMN amplitudes. A range of other studies conducted in languages with a fixed stress pattern, such as Hungarian (Honbolygó & Csépe, 2013) or Finnish (Ylinen et al., 2009) does however show a modulation that is in line with a violation response (i.e. larger MMN to the illegal stress pattern). The divergent results here indicate that Dutch speakers process variations in lexical stress patterns differently than their Hungarian or Finnish speaking counterparts. While syllable stress may be exploited during development (Weber et al., 2004), and continues to play a role in resolving lexical conflict in spoken word recognition (Cutler & Van Donselaar, 2001), predictions generated relating to stress patterns may be weaker compared to those relating to phonotactic probability, particularly in the case of pseudoword stimuli in languages without a fixed lexical stress pattern.

### P2

We further investigated the effect of variations of phonotactic probability and syllable stress across self- and externally generated conditions on P2 mean amplitude. Here, we observe a main effect of syllable stress, where first syllable stress stimuli elicit a larger P2 amplitude compared to second syllable stress stimuli. This observation may be explained by acoustic differences in first and second syllable stress items (see Figure 1). The main acoustic markers of lexical stress are intensity, pitch, and duration of the syllable. Thus, while the stimuli were equalized in intensity across the whole word, they differed in the first syllable, with first syllable stress items having a greater intensity compared to second syllable stress items. The timing of our P2 at around 280 ms suggests that this component reflects information from the first syllable. We do not observe an interaction with condition, which would suggest a conscious differentiation between self- and externally generated events (Knolle et al., 2013a, 2019; Pinheiro et al., 2018). Therefore, this pattern is more in line with previous observations of P2 amplitude being modulated by stimulus intensity (for review, see Crowley & Colrain, 2004). However, if this is indeed a purely acoustic effect, one would expect to find a similar modulation in the N1 component, which we do not observe. It is possible that the amplitude modulation from the syllable stress pattern is masked due to distortion of the overall N1 amplitude from overlap of the pre-stimulus deflection. However, assuming that this distortion is equal across stimuli we would still expect to observe an effect in the N1 amplitude across first and second syllable stress stimuli.

An alternative explanation for the P2 modulation may lie in categorical perception of speech. The neural correlates of categorical perception around the typical P2 time-window, indicated by investigations of phoneme processing comparing tokens that vary along a continuum across phoneme boundaries (e.g. Bidelman et al., 2013, 2020). Here, ambiguous speech sounds show a smaller P2 amplitude compared to speech sounds that clearly fall within a phoneme category. As second syllable stress is atypical for Dutch bisyllabic words, this may come with variability in the pronunciation, including variability in vowel quality. However, as the current experiment did not specifically modulate categorization, and we also do not have data on the perceptual categorization of the vowels in the current design, the current data cannot address this question completely, thus this interpretation remains speculative.

### General discussion

The temporal dissociation of the observed effects, with phonotactic probability modulating N1 and syllable stress P2 amplitude, may suggest differences in the time course of processing of these features. In the current design, the features phonotactic probability and syllable stress are manipulated at different points in time of the stimuli: first and second syllable stress items differ from each other in principle from stimulus onset on, while phonotactic probability is varied at the syllable boundary. Furthermore, as the stimuli are naturally produced, we have little control over the precise timing of acoustic markers of phonotactic probability and syllable stress relative to the timing of the ERP components of interest. This makes it difficult to disentangle differences in the time course of the neural processing of these linguistic features from differences in when the information relevant to these features becomes available in the specific pseudoword stimuli. Thus, it is not possible to draw general conclusions about the relative time course of processing of phonotactic probability and syllable stress beyond the current design.

Although we originally set out to test hypotheses relating to motor-induced suppression, limitations to the current design hinder us from following the original analysis plan. The pre-stimulus deflection observed in the auditory-only condition (Figure 3) draws attention to the cue as a confound. This deflection appears to be time-locked to the cue onset, covers a broad time window and is present across the scalp, though larger in amplitude at more parietal regions. While the cue is identical in all conditions, it is effectively subtracted out from the motor-auditory condition along with the motor component (MAC = MA - MO) but remains present in the auditory-only condition. Including a visual control to subtract from the auditory-only condition may ameliorate this issue, however this would only account for purely visual processes. The deflection likely also represents attentional and anticipatory processes, as the participant was instructed to explicitly attend to the stimulus and could anticipate not only *which item* would be presented, but also *when* it would be presented, due to the constant timing between cue and stimulus. Thus, an additional adjustment to the current paradigm could include jittering the timing of these events to dissociate the processes associated with the cue and the stimulus. Varying the time between cue and stimulus could also address the question of whether the suppression effect is driven by the temporal predictability of the stimulus (Hughes et al., 2013; Sowman et al., 2012).

We do not observe any suppression in either the N1 or P2 components for the motor-auditory vs auditory-only condition. This observation differs from the bulk of similar studies applying this type of paradigm comparing the processing of self- and externally-generated stimuli (e.g., Bäss et al., 2008; Knolle et al., 2012, 2013a; Pinheiro et al., 2018). The lack of suppression may be explained by variations in the design between previous studies and the current experiment. The typical approach in this paradigm does not include a cue. Instead, the paradigm is typically applied as a blocked design (but see Knolle et al., 2013b for an event-related variation), where the button-presses generating the stimulus presentation are self-initiated in the motor-auditory condition. The auditory stimuli are then presented at the same temporal intervals in the auditory-only condition, again without a cue. Thus, a crucial difference between the auditory-only and motor-auditory conditions in these approaches, in addition to whether the stimulus is self-generated or externally presented, is the predictability of the stimulus timing: in the motor-auditory condition, the participant can accurately predict the timing, while some temporal uncertainty remains in the auditory-only condition. A considerable portion of the suppression effect observed in previous research may therefore be driven by the temporal predictability of the events. In the current study, the timing of the stimulus in the auditory-only condition is predictable due to the cue, thus this difference between the auditory-only and motor-auditory conditions does not exist. If temporal predictability indeed drives the suppression effect, it is therefore not surprising that we do not observe this effect in the current paradigm. Future studies investigating the suppression effect should therefore consider not only varying the formal predictability of the stimulus, but also its temporal predictability.

In conclusion, the present experiment provides preliminary insights into differences in processing phonotactic and temporal regularities in speech perception and production, by comparing self-generated (via button press) to externally generated (own) speech. Our findings suggest that phonotactic regularities play an important role in processing self-generated speech, with a perceptual bias toward more probable phonotactic structures, in line with Bayesian accounts of predictive processing, or a combined model incorporating both Bayesian and cancellation theories. We further observe an effect of syllable stress, which is likely explained by the acoustic differences between the first syllable in pseudowords with first and second syllable stress. To summarize, the current research suggests that a sensitivity to regularities in phonotactic and temporal structure of speech may be differently exploited in speech perception and production processes. Further investigations controlling for some of the limitations observed in the current paradigm are needed to confirm the results of the current post-hoc analyses.

## Data availability

The data that support the findings of this study are available from the corresponding author upon reasonable request.

## Acknowledgements

The authors would like to thank Joseph Johnson and Stevan Nikolin for sharing code that was adapted for the current experiment, Dana Hermsen and Alexa Holfelder for their assistance with data collection, and Suvarnalata Xanthate Duggirala, Francesco Gentile, Lisa Goller, Lars Hausfeld and Michael Schwartze for discussions on data analysis. This research was supported by Maastricht University (Incentives to BMJ for deanship at Faculty of Science and Engineering) and The Netherlands Organization for Scientific Research (Vidi-Grant 452-16-004 to MB).

## Author Contributions

AKE, MB, BMJ and SAK designed the experiment, AKE prepared materials, collected and analyzed the data, and wrote the first draft of the manuscript, AKE, MB, BMJ and SAK refined the manuscript.

## Competing Interests

The authors declare that they have no competing interests.

### Abbreviations

AO: auditory-only,
BOLD: blood oxygenation level dependent,
Cond: condition,
ERP: event-related potential,
fMRI: functional magnetic resonance imaging,
HPP: high phonotactic probability,
ICA: independent component analysis,
IQR: inter-quartile range,
LPP: low phonotactic probability,
MA: motor-auditory,
MAC: motor-auditory corrected,
M/EEG: magneto-/electroencephalography,
MIS: motor-induced suppression,
MO: motor-only,
PhonProb: phonotactic probability,
ROI: region of interest,
SylStr: syllable stress,
SylS1: first syllable stress,
SylS2: second syllable stress.

## Notes

### Competing Interest Statement

The authors have declared no competing interest.

