## Supplementary Materials for "Phonological and temporal regularities lead to differential ERP effects in self- and externally generated speech"

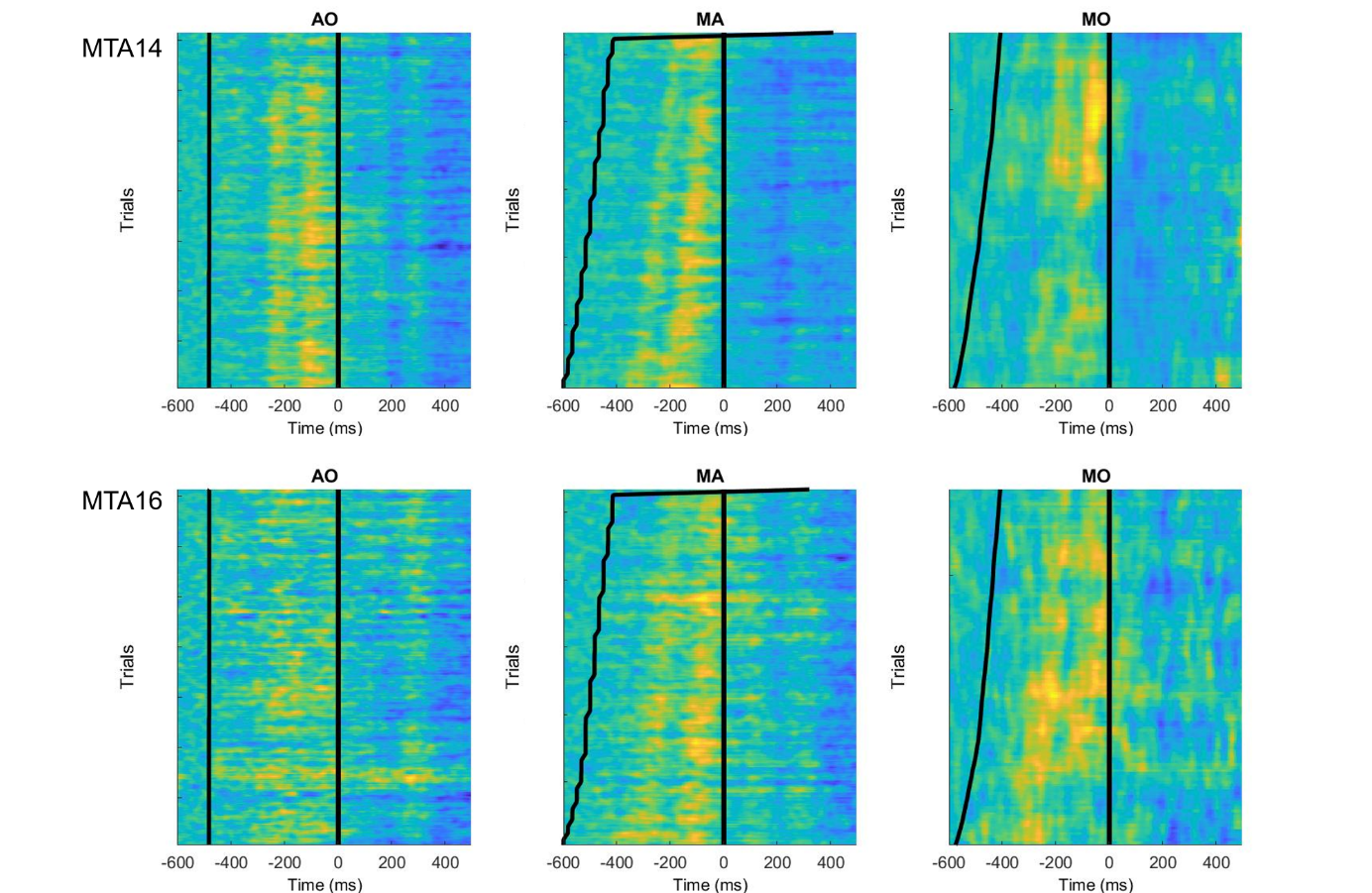

Figure S1: Exemplary participants MTA14 and MTA16 illustrating pre-stimulus deflection is related to visual cue. Single trial data at channel FCz represented on the y-axis, color representing amplitude in arbitrary scale (yellow = positive, blue = negative). In MA and MO conditions, trials are sorted based on interval between cue (black line at around t = -500 ms) and stimulus (black line at t = 0). Pre-stimulus positivity is aligned with timing of cue in both MA and MO condition.

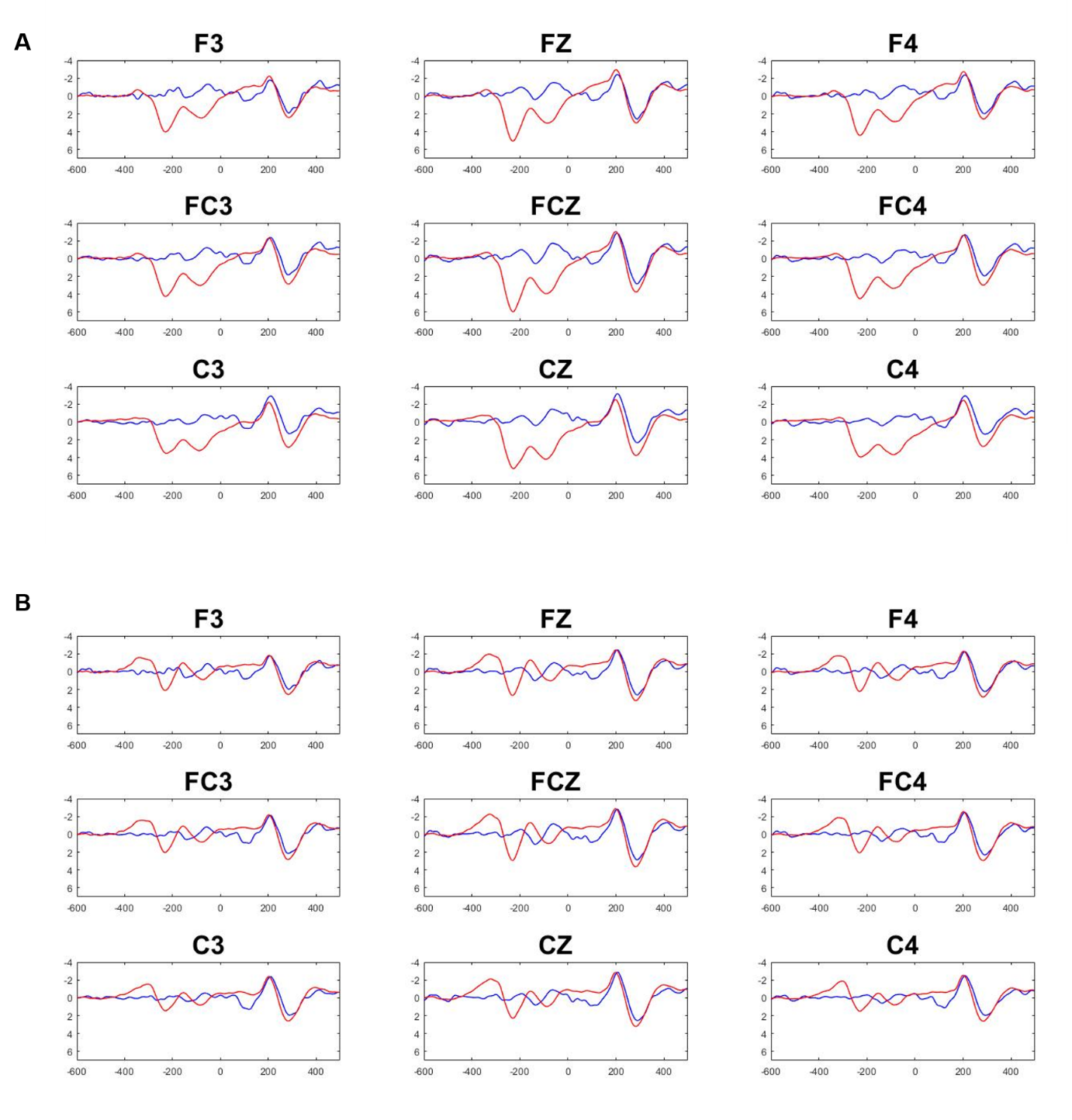

Figure S2: AO vs MAC -600 to 500 ms relative to stimulus onset in sample of n = 22. (A) 1-30Hz filter used in final analyses (B) 3-30Hz filter. Higher high-pass cutoff reduces overall amplitude of pre-stimulus deflection but does not remove it.

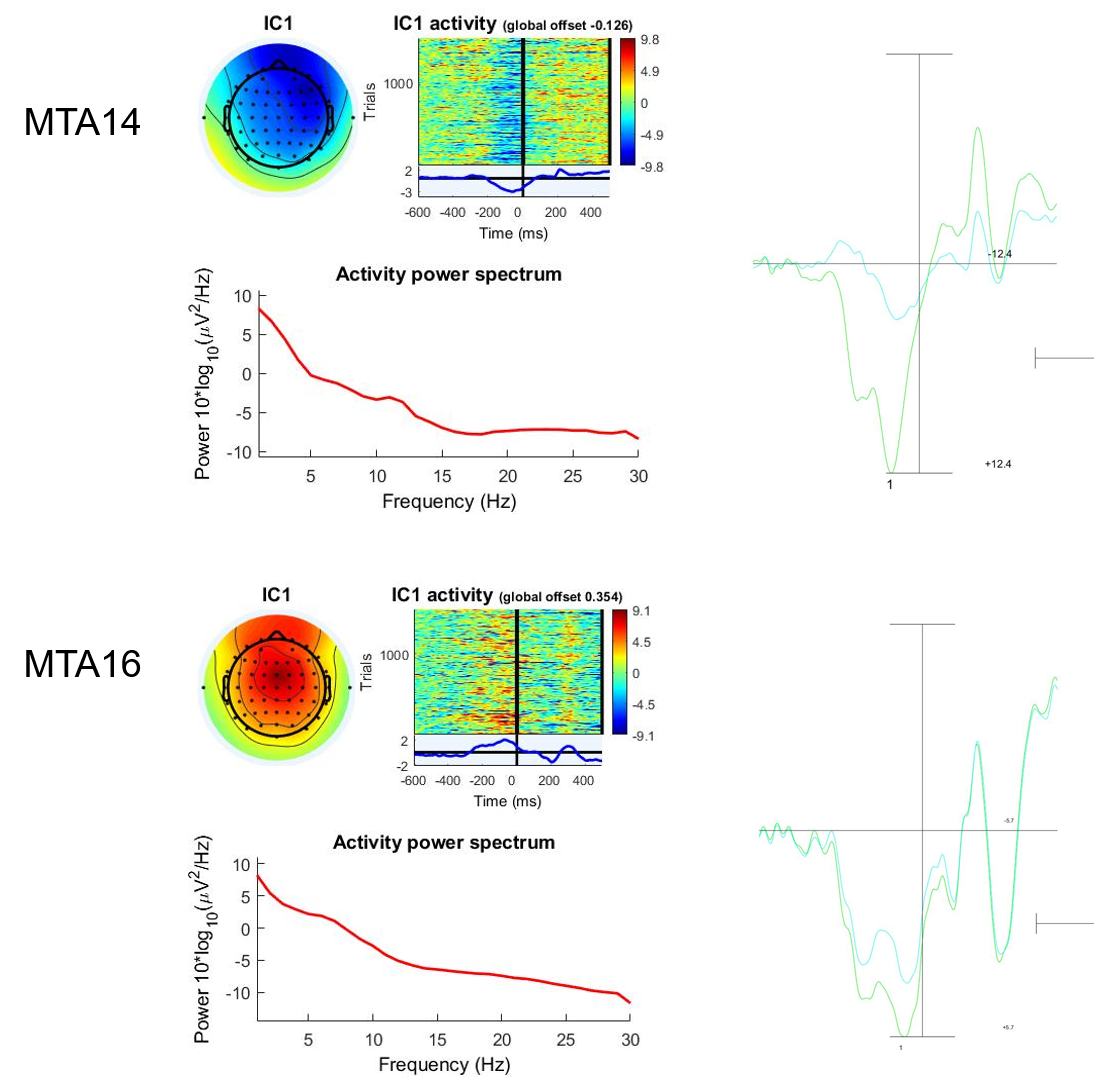

Figure S3: Exemplary participants MTA14 and MTA16: ICA fails to separate pre-stimulus deflection from components of interest. Left: topography, time course and power spectrum of independent component (IC) containing pre-stimulus deflection. The same component also includes deflection in the time window of N1 and P2 effects. Right: grand average ERP at channel FCz across all conditions. Green includes all ICs, blue with the selected IC removed. Removing this component often substantially reduces N100 amplitude (MTA14) or does not affect the positivity as desired (MTA16).

Supplementary Table S1

*Descriptive statistics N1 mean amplitudes*

| **Within-subjects variables** | | | **N1 mean amplitude** | | **Shapiro-Wilk test** | |
| --- | --- | --- | --- | --- | --- | --- |
| **Cond** | **PhonProb** | **SylStr** | **Mean** | **Std Dev** | **W** | **p** |
| **AO** | **HPP** | **SylS1** | -2.94 | 2.48 | 0.967 | 0.536 |
|  |  | **SylS2** | -2.80 | 2.58 | 0.983 | 0.919 |
|  | **LPP** | **SylS1** | -2.67 | 2.75 | 0.934 | 0.087 |
|  |  | **SylS2** | -3.44 | 2.52 | 0.957 | 0.319 |
| **MAC** | **HPP** | **SylS1** | -3.32 | 2.01 | 0.980 | 0.853 |
|  |  | **SylS2** | -3.02 | 1.82 | 0.957 | 0.314 |
|  | **LPP** | **SylS1** | -2.68 | 2.66 | 0.981 | 0.891 |
|  |  | **SylS2** | -2.67 | 2.51 | 0.976 | 0.763 |

Supplementary Table S2

2x2x2 Repeated measures ANOVA on N1 mean amplitude (Bonferroni-Holm corrected)

| **Effect** | **F(1,26)** | **p_adj_** | **Effect size η^2^_p_** |
| --- | --- | --- | --- |
| **PhonProb** | 0.605 | 1.000 | 0.023 |
| **SylStr** | 0.238 | 1.000 | 0.009 |
| **Cond** | 0.010 | 1.000 | <0.001 |
| **PhonProb x SylStr** | 4.222 | 0.300 | 0.140 |
| **PhonProb x Cond** | 8.463 | 0.049 * | 0.246 |
| **SylStr x Cond** | 3.541 | 0.355 | 0.120 |
| **PhonProb x SylStr x Cond** | 2.624 | 0.468 | 0.092 |

Supplementary Table S3

Post-hoc paired samples t-tests of effect of PhonProb at individual levels of Condition. (HPP vs LPP)

| **Condition** | **t(26)** | **p_adj_** | **Effect size (d)** |
| --- | --- | --- | --- |
| **AO** | 0.840 | 0.818 | 0.162 |
| **MAC** | -2.04 | 0.104 | -0.392 |

Supplementary Table S4

*Descriptive statistics P2 mean amplitudes*

| **Within-subjects variables** | | | **N1 mean amplitude** | | **Shapiro-Wilk test** | |
| --- | --- | --- | --- | --- | --- | --- |
| **Cond** | **PhonProb** | **SylStr** | **Mean** | **Std Dev** | **W** | **p** |
| **AO** | **HPP** | **SylS1** | 3.56 | 3.42 | 0.988 | 0.984 |
|  |  | **SylS2** | 2.70 | 3.00 | 0.973 | 0.688 |
|  | **LPP** | **SylS1** | 3.46 | 3.35 | 0.961 | 0.387 |
|  |  | **SylS2** | 2.37 | 3.29 | 0.979 | 0.834 |
| **MAC** | **HPP** | **SylS1** | 3.05 | 2.60 | 0.966 | 0.493 |
|  |  | **SylS2** | 1.96 | 2.18 | 0.979 | 0.828 |
|  | **LPP** | **SylS1** | 3.14 | 2.69 | 0.975 | 0.741 |
|  |  | **SylS2** | 2.53 | 2.43 | 0.970 | 0.596 |

Supplementary Table S5

2x2x2 Repeated measures ANOVA on P2 mean amplitude (Bonferroni-Holm corrected)

| **Effect** | **F(1,26)** | **p_adj_** | **Effect size η^2^_p_** |
| --- | --- | --- | --- |
| **PhonProb** | 0.065 | 1.000 | 0.003 |
| **SylStr** | 22.993 | 0.0004*** | 0.469 |
| **Cond** | 0.760 | 1.000 | 0.028 |
| **PhonProb x SylStr** | 0.092 | 1.000 | 0.004 |
| **PhonProb x Cond** | 4.292 | 0.288 | 0.142 |
| **SylStr x Cond** | 0.232 | 1.000 | 0.009 |
| **PhonProb x SylStr x Cond** | 1.508 | 1.000 | 0.055 |
